# TUCL-Seq for miRNA single molecule sequencing

**DOI:** 10.1101/2025.04.07.646966

**Authors:** Jicai Fan, Zhao Li, Xuejiao You, Lei Liu, Huan Jin, Ping Wu, Yushan Han, Lei Sun, Pengwei Xu, Yongfeng Liu, Jianjun Luo, Qin Yan

**Author notes:** For correspondence: Qin Yan. Address: 5F and 6F, Block 2, Luohu Investment Holding Building, No. 116, Qingshuihe 1st Road, Luohu District, Shenzhen, Guangdong, 518000, China.

## Abstract

MicroRNAs (miRNAs) are pivotal regulatory molecules in gene expression, and their precise quantification is essential for elucidating their roles in diverse biological processes. This research paper introduces a pioneering workflow for the single molecule sequencing (SMS) of miRNA, integrating advanced miRNA sample preparation with SMS sequencing. Central to our approach is TUCL-Seq, which is an automated, on-chip miRNA library preparation method designed to streamline the sequencing process. This innovative approach incorporates the addition of sequencing adaptor to RNA samples in solution, and converts it to cDNA on a sequencing flow cell for sequencing. Coupled with single molecule sequencing instrument and base-call software, our workflow significantly reduces sample-to-result time, and decreases the need for manual interventions compared to traditional SMS and next-generation sequencing methods. We demonstrate the efficacy of TUCL-Seq by accurately quantifying synthetic miRNA samples with a 24-hour workflow, showcasing its potential for high-throughput miRNA profiling. Our findings highlight the promise of this integrated workflow for advancing miRNA research and applications in diagnostics and therapeutics.

## Introduction

MicroRNAs (miRNAs) have emerged as an important biomarker, owing to their intricate regulatory roles in gene expression and their implications in various physiological and pathological processes. Traditionally, next-generation sequencing (NGS) is a powerful tool for miRNA profiling (1). However, existing NGS methods, which involve converting miRNAs to cDNA through reverse transcription and subsequent amplification, typically require lengthy sample preparation and are prone to amplification and ligation biases (2,3).

The introduction of single molecule sequencing (SMS) has revolutionized the way for RNA research, providing unique advantages over NGS methods by eliminating amplification and enhancing quantification accuracy. However, the full potential of SMS in miRNA analysis has been hindered by laborious and error-prone library preparation protocols. The low abundance, sequence similarities, and short length of miRNAs also create considerable difficulty for long-read SMS technologies, including single-molecule real-time (SMRT) and nanopore sequencing (4). Recently published nanopore sequencing of miRNA has shown the identification of different sequences, isoforms, and epigenetic modifications among synthetic miRNA sequences (5), however, accurate detection remains challenging due to nanopore’s limited throughput and sample amount requirement. Consequently, quantitation of miRNA has not been attempted.

Another SMS method available is pioneered by Helicos (6), which utilized a visual terminator and total internal reflection fluorescent microscope (TIRF). Direct RNA sequencing was reported using this approach (7), showing the quantitation of mRNAs by capturing the poly-A tails and sequencing after the “fill lock” procedure. Recent research on sequencing platform GenoCare 1600 introduced two-color sequencing chemistry to this SMS method, leading to new applications in microbial sequencing (8) and non-invasive prenatal testing (9). Though GenoCare 1600 generates a lower read length (50-70 nt) compared to the other two SMS methods, its higher reads throughput and simpler library preparation could be exploited for miRNA sequencing. Since miRNAs are usually between 18 and 30 nt long (10), the read length of GenoCare 1600 is sufficient to characterize them.

Introducing sequencing adaptor oligodeoxyribonucleic acid (ODN) to miRNA is an important step before sequencing. Bullard reported that T4 RNA ligase could efficiently link the 5’ end of the donor ODN to the 3’ end of the recipient RNA (11), while, it is difficult to attach the 3’ end of the DNA. The length of the tail with homopolymer tail added to the DNA by terminal deoxyribonucleotidyl transferase (TdT) depends on the nucleotide types used, and it is difficult to control the length of the added tail. When using NTPs, the tail length is obviously self-limiting, as the reaction is automatically terminates after adding 2∼4 nt with a high efficiency, above 98% (12). Besides, Miura used the characteristics of T4 RNA ligase and TdT to develop an efficient ssDNA ligation method (TACS, **T**dT-assisted **a**denylate **c**onnector-mediated **s**sDNA ligation). In brief, TdT was used to add adenosine to the 3’ end of the ssDNA, and then T4 RNA ligase was used to link the 5’ end of the ODN to the ssDNA after the addition of adenylate to the 3’ end (13).

This research paper provides automated, on-chip library preparation to streamline the single molecule sequencing process (TUCL-Seq, **t**erminal deoxyribonucleotidyl transferase-assisted **u**ridine **c**onnector-mediated 1st cDNA **l**igation **seq**uencing) for miRNA analysis. We delve into the methods of integrated library preparation of miRNAs from synthetic samples, and a novel single molecule sequencing scheme of the resulting libraries. We also scrutinize the challenges associated with different sequencing methods and optimize the base-calling and bioinformatics algorithms.

## Results

### Design of TUCL-Seq for miRNA

We developed TUCL-Seq, a method based on the TACS approach, streamlined manual steps in the library construction process and performed automated integration (Figure 1). Initially, a poly(A) tail is added to 3’ end of the miRNAs using PAPase. The high conversion rate makes purification unnecessary, allowing direct capture by poly(dT50LNA2)probes on the SMS flow cell. Two LNA modifications were introduced near the 3’ end of the poly(dT50) probe to enhance the capture efficiency and stability (Figure S1). On the flow cell surface, reverse transcriptase converts RNA targets to cDNAs, and excess poly(dT50 LNA2) probes are digested by Exo I to prevent interference in subsequent reactions. The miRNA/cDNA double strand is then denatured, and the miRNA template is eluted. Next, TdT adds 2∼4 rU to the 3’ end of the resulting 1^st^ cDNA, and proteinase K removes the TdT adsorbed on the chip surface. T4 RNA ligase 1 subsequently links the 5’ end of the ssDNA sequencing adapter to the 3’ end of the 1^st^ cDNA. The sequencing primer is hybridized, and polymerase with dATP is introduced to fill in the single-stranded polyU and polyT regions. Sequencing then proceeds. The entire process, from RNA hybridization capture to pre-sequencing, is integrated into a liquid handling instrument named X-bot.

**Figure 1.**
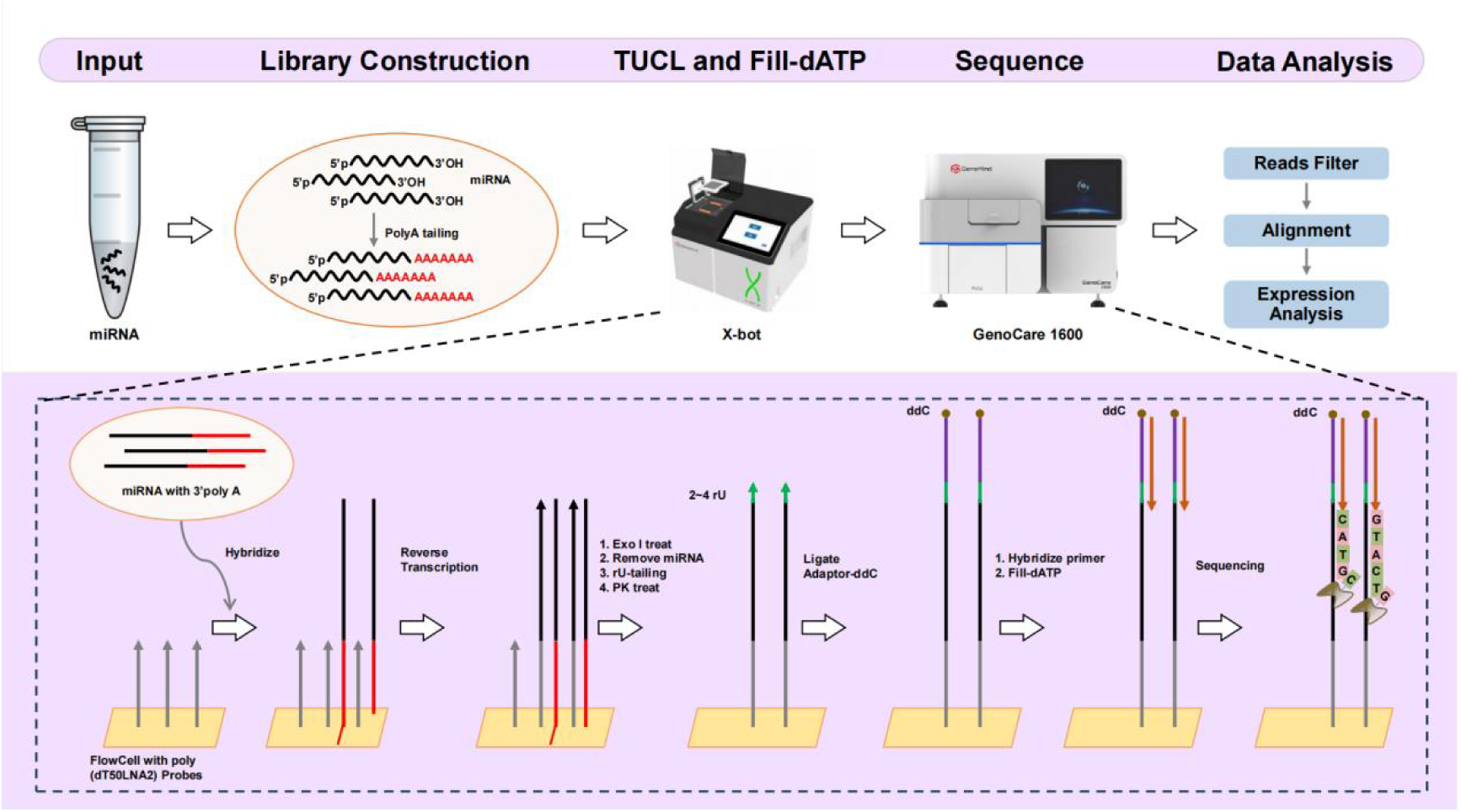
A diagram depicting the TUCL-Seq workflow.

### rU-tailing and 3’ bias Optimization

Previous report has indicated that TdT utilizes NTPs to add homopolymer tails to ssDNA ends with self-limiting length (12). However, the molar ratio relationship between NTPs and ssDNA and the potential bias introduced by different 3’ bases on ssDNA have not been investigated. To address this, we designed five ssDNA strands with varying 3’ bases (Table 1) to examine the stoichiometry relationship between rU-tailing, NTPs, and ssDNA, as well as the influence of the 3’ bases on ssDNA. Our findings revealed that when the UTP to ssDNA ratio was ≥ 10:1, 3-4 rU residues could be consistently added to nearly all ssDNA ends (Figure S2A). Moreover, we observed no rU-tailing bias across different 3’ ends, with the efficiency approaching 100% (Figure S2B). The rU-tailing efficiency on the SMS flow cell surface was similarly high, with an efficiency close to 100% (Figure S2C).

**Table 1.**
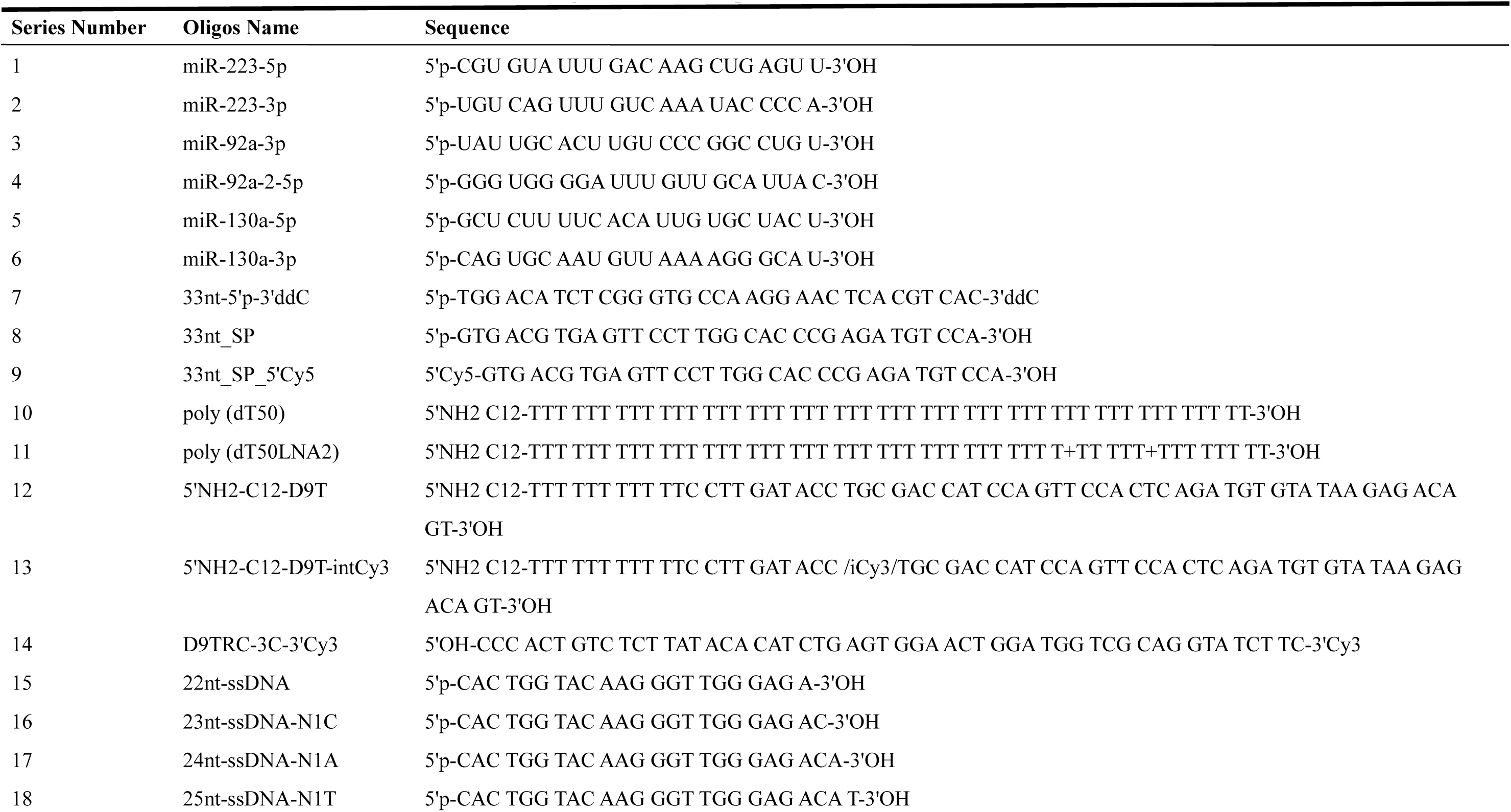
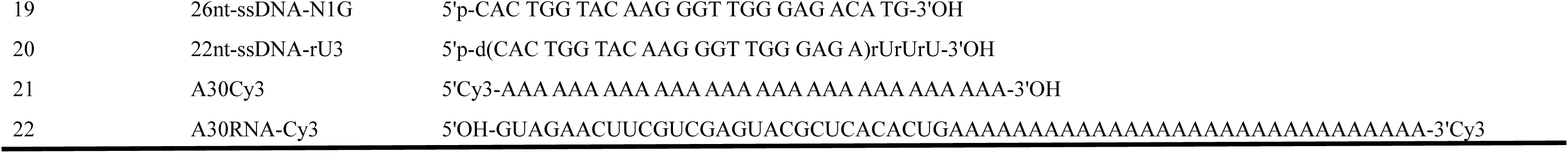
Oligos and their sequences used for ssDNA strands.

### ssDNA Ligate Optimization

Although the TACS method enhances the ligation efficiency of ssDNAs, its conversion rate remains at 70% (13). RNA ligase catalyzes the RNA-DNA ligation reaction in a three-step process (14): first, it reacts with ATP to form an adenylyl intermediate, releasing pyrophosphate; next, the adenylyl intermediate transfers its adenylate (AMP) group to the 5’ phosphate of ssRNA or ssDNA, creating an activated 5’ adenylyl intermediate (5’AppRNA or 5’AppDNA); finally, RNA ligase facilitates the substitution of the 3’-OH of the acceptor substrate for the 5’-AMP of the donor, forming a phosphodiester bond and completing the ligation (15–17) We discovered that with rU-tailed ssDNA, the ligation efficiency in solution can reach up to 90% when the molar ratio of ATP:ssDNA:RNA ligase is 5:1:1∼50:1:1 (Figure 2A).

**Figure 2.**
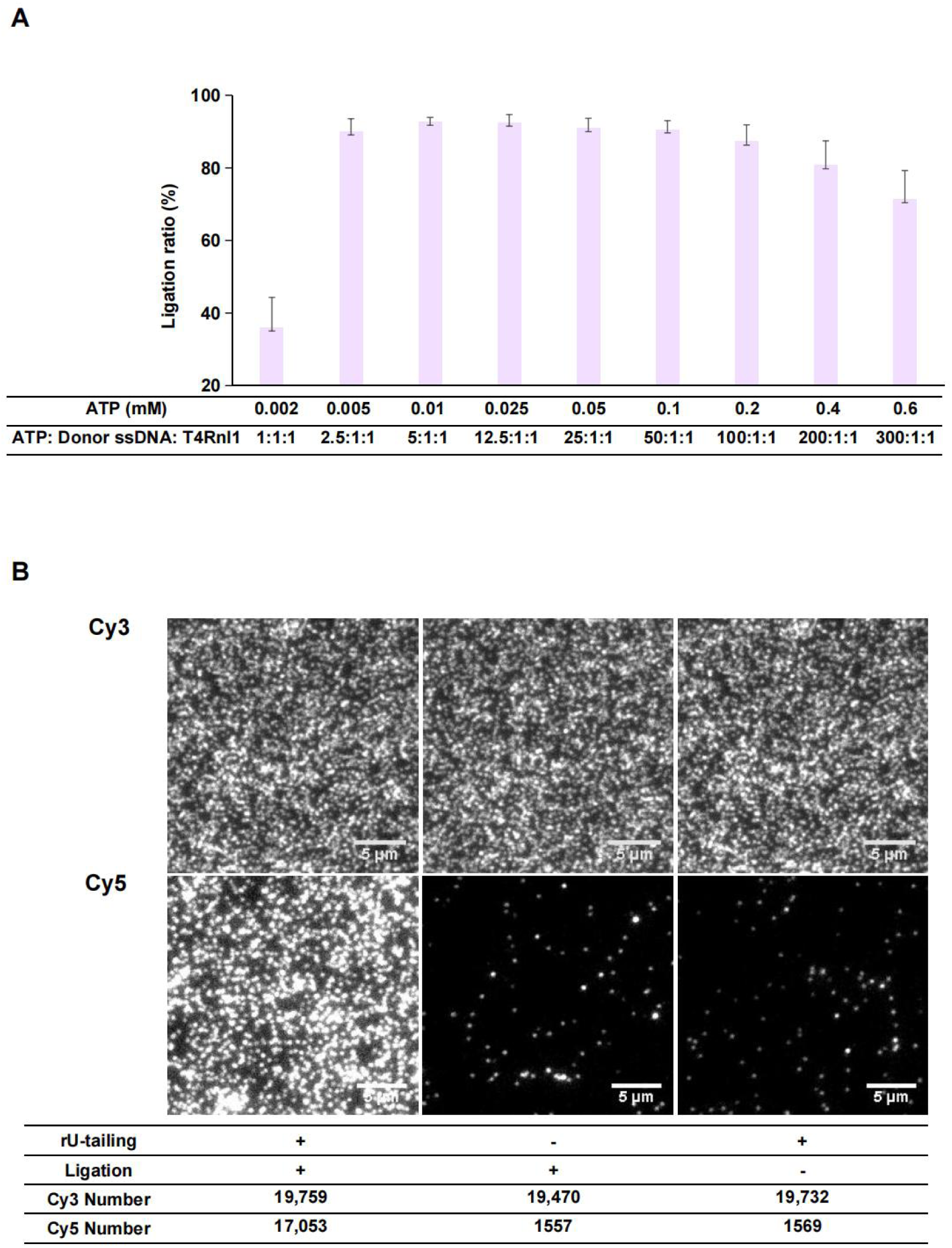
The efficiency of ssDNA ligation. A) Assessment of the ligation efficiency between the 3’ uracilated ssDNA acceptor and the 5’ phosphorylated ssDNA donor, catalyzed by T4 RNA Ligase 1, at varying ATP concentrations. B) Evaluation of the ligation efficiency of TUCL on the surface of the SMS flow cell. The acceptor ssDNA is labeled with Cy3, and the donor ssDNA is labeled with Cy5. The accompanying table presents the fluorescence spot count within a 1k×1k pixel image field of view.

Furthermore, testing the reaction on the SMS flow cell surface, we achieved a comprehensive efficiency of >85% in ligating surface-bound and rU-tailed ssDNA to the 33nt-5’p-ddC adapter (Figure 2B, the adaptor’s 3’ end is blocked with a dideoxy cytosine base to prevent side reaction). In contrast, ssDNA lacking rU-tailing or treated without ligase produced minimal fluorescence signals.

### Surface Superfluity Probes Removal

Unreacted surface primers that do not participate in the first cDNA generation could engage in rU-tailing and subsequent ligation, leading to significant sequencing noise. To circumvent this, we digested these primers with Exo I after reverse transcription and before denaturing the eluted RNA template. Since TdT’s non-template-dependent polymerase activity requires a primer with at least three bases, Exo I-treated ssDNA cannot be recognized by TdT. We investigated the excision efficiency of poly (dT50LNA2) single strands on the SMS flow cell surface by Exo I and achieved a resection rate of 96.01% after a 15 min treatment at 37℃. Further treatments did not enhance the outcome (Figure S3B&C). In parallel studies, we used Exo I to remove D9T single strands from the SMS flow cell surface and observed a slower digestion rate (Figure S3A). We hypothesize that most ssDNA on the flow cell surface is readily excisable, while a small portion absorbs onto the surface and resists digestion. This absorption may result from interactions between the amino groups of the ssDNA bases and the epoxy groups on the SMS flow cell surface during surface immobilization. The poly (dT50LNA2)LNA2 surface primer can be removed more effectively even with the LNA base modifications, as it lacks such amino groups in its base structure.

### Performance of six miRNAs Mixture Quantification

To evaluate the quantitative capabilities of TUCL-Seq for miRNA detection, we synthesized six miRNAs related to colorectal cancer. These RNAs were mixed in a ratio of 1:10:100:1000:10000:100000, and the starting amounts for different library preparations were set to 1, 0.5, 0.1, and 0.05 pmol for TUCL-Seq. Each experiment was conducted in quadruplicate. Our findings indicate that when the initial miRNA amount ranged from 0.1 to 1 pmol, the quantitative results were relatively accurate across four orders of magnitude, from 100 to 1,000,000 (counts per million reads, CPM). The corresponding detection limit of miRNA is roughly at 0.1 fmol (0.7 pg) when initial miRNA amount is 0.1 pmol. The quantitative accuracy diminished when target miRNA amount is lower, or when the initial miRNA amount was less than 0.1 pmol (Figure 3).

**Figure 3.**
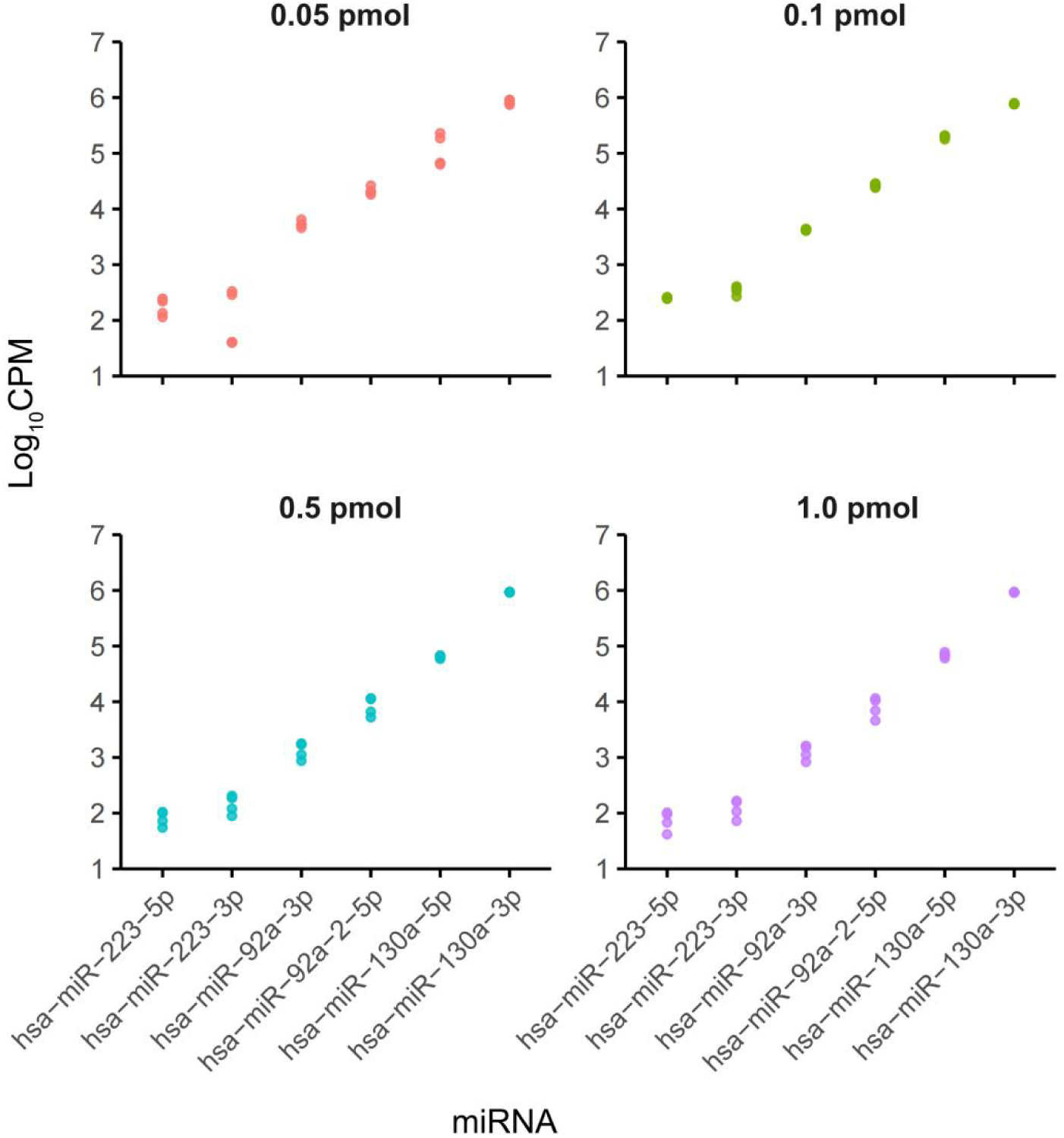
Scatter plot of miRNA quantification at different concentrations. CPM means counts per million reads.

## Discussion

Since miRNA does not have a poly-A tail, prior reported directing single-molecule RNA sequencing method is incompatible. Usual protocols for construction of miRNA library can be broadly summarized into two categories: (A) ligation of 3- and 5-terminal adaptors by ligase, first by ligation of an ssDNA adapter at the 3’ end by T4 RNA ligase, and then by purification or digestion to remove the excess 3-terminal adaptor, followed by T4 RNA ligase attaches an RNA adapter to its 5’ end, and finally the cDNA library is obtained by RT-PCR (18); (B) The 3- and 5-terminal adaptors were added by SMARTer technology, firstly, a poly(A) tail was added to the 3’ end of PAPase, then a TSO (template switch oligo) adapter was added to the 3’ end of the first-strand cDNA while reverse transcription was obtained by template conversion activity of reverse transcriptase, and finally the cDNA library was obtained by PCR amplification (19). Scheme A is rather long and labor-intensive; while scheme B has low template conversion efficiency due to the absence of a cap structure at the 5’ end of the miRNA, and there are serious 5’ base bias and multi-template conversion problems (20).

To develop the on-surface miRNA library construction method, we first explored the less complicate Scheme B and tried to developed a surface-based method. However, while ligating an A-tail to the miRNA and then generating cDNA on a flow cell using A-tail capture proceeded smoothly, The following steps to add 3’-adaptors to cDNA proved to be complex and the yield was very low, partly due to the inefficient TSO reaction on surface. TACS method could improve the efficiency of cDNA conversion in solution, but we found it does not perform as well on flow cell surface. In contrast, the TUCL-Seq method allowed efficient addition of sequencing adaptors to the cDNAs generated on surface, and is applicable to a variety of RNA targets. Theoretically, this method could be applied to NGS sequencing platform as well if one of the surface amplification primers could be modified to include a polyT region.

The automated library preparation platform streamlined the workflow and expedited the miRNA sequencing. By integrating steps including adapter ligation, reverse transcription, and sequencing primer hybridization, this method minimizes hands-on time, reduces sample input requirements, and lowers the risk of sample contamination. The 2-color sequencing chemistry also significantly reduced sequencing time compared with the prior art. Starting from extracted miRNA, the entire process could be completed within 24 hours. While, NGS is very time consuming and still is not fully automated, with the turnaround time of about four days (21,22). The RNA degradation issue is also mitigated, as we converted miRNA to the more stable cDNA for the later sequencing steps.

In previous single-molecule RNA sequencing works (7), low accuracy of the single molecule method proved to be an issue that prevented accurate quantitation of miRNAs. Here we built the basecall method based on the Smith-Waterman algorithm, modifying minimum read length and error base thresholds and optimizing the parameter using simulated sequencing data (supporting information: method section). We found that the condition of read length >15 nt and maximum error rate of 10% yielded the best throughput-accuracy combination. Considering the real RNA sequencing data could contain a large portion of “noise reads”, including adaptor sequences and signals from non-specific binding from the SMS platform, we tried to construct a “noise sequence” database and utilize it as a background. However, we found that these noises did not impact the quantitation accuracy as much when we optimized the data filtering threshold, and the noise sequence database did not improve the quantitation results (data not shown).

Despite the advantages of this single molecule RNA sequencing method, this approach is limited by several factors: firstly, the accuracy of SMS is still below the NGS method. As a result, it is more suitable for small panels of miRNA. Secondly, since no amplification is performed, the single molecule method requires a relatively large amount of sample. Lastly, this method hasn’t been tested on clinical samples due to the large amount of miRNA required. Future study would be focused on reducing the starting sample amount and improving the sequencing accuracy.

In this study, we introduced a comprehensive workflow that integrates miRNA analysis with SMS results. This workflow features an automated, on-chip miRNA library preparation technique called TUCL-Seq, along with a SMS instrument and associated software. Compared to traditional SMS and NGS methods, TUCL-Seq exhibits reduced bias and a shorter sample-to-result time. Additionally, we demonstrated the capability to quantify synthetic miRNA using this approach. We aim for this research to stimulate further advancements in the field, enhance our understanding of miRNA biology, and open avenues for novel clinical applications.

## Experimental procedures

### miRNA and oligo DNA synthesis

The six miRNAs associated with colorectal cancer (23,24) along with the oligonucleotides 33nt-5’rApp-ddC and 59nt_A30RNA-Cy3, were procured from BiOligo Biotechnology (Shanghai) Co., Ltd. Additional oligonucleotides listed in Table 1 were sourced from Sangon Biotech (Shanghai) Co., Ltd.

### Flow Cells and Surface Chemistry

The SMS sequencing chip and its surface chemical modification are processed as described previously (25). To enhance the on-chip hybridization stability between the miRNA 3’ poly(A) tail and the poly (dT50) probe during reverse transcription, we designed two locked nucleic acid modifications near the 3’ end of the poly (dT50) probe, as poly (dT50LNA2) in Table1.

### TUCL-Seq Library Preparation and Sequencing

Six miRNAs were dissolved in NF-H2O (Invitrogen, cat. No. 10977015) to a concentration of 40 μM, and then diluted in a 10-fold gradient: miR-223-5p to 100,000 times, miR-223-3p to 10,000 times, miR-92a-3p to 1,000 times, miR-92a-2-5p to 100 times, miR-130a-5p to 10 times, and miR-130a-3p undiluted. Equal volumes of each dilution were mixed to create the miR6mix library.

For the preparation of 1 pmol miR6mix, add NF-H2O to a final volume of 7.25 μl, then mix with 1 μl of 10× poly(A) polymerase reaction buffer (500 mM Tris-HCl, 2500 mM NaCl, 100 mM MgCl2, pH 8.1 @ 25°C), 1 μl of ATP (diluted to 0.2 mM, NEB, cat. No. P0756S), 0.25 μl of Murine RNase Inhibitor (Vazyme Bio, cat.No. R301-03), and 0.5 μl of *E. coli* poly(A) polymerase (NEB, cat. No. M0276L). Incubate at 37℃ for 30 minutes. After incubation, add 0.3 μl of 0.5 M EDTA to terminate the reaction and place on ice. The protocol can be applied to RNA quantities below the recommended 1 pmol level (down to 0.05 pmol), with adjustments to the ATP concentration to maintain a ATP:miRNA ratio 200:1.

The miR6mix library was diluted to 10 nM with NF-H2O. Mix 1.6 μl of this solution with 13.4 μl of NF-H2O and 25 μl of hybridization reagent V2 (Genemind Biosciences Co., Ltd, Shenzhen, China). Load the hybridization mixture onto a GenoCare flow cell using the X-bot automated liquid handler (Genemind Biosciences Co., Ltd, Shenzhen, China). The GenoCare flow cell has 16 lanes, with each lane capable of testing one sample and requiring 40 μl of the mixture. Incubate the flow cell at 37℃ for 20 minutes, then wash away the unhybridized libraries. The detailed steps are as follows.

| Step | Description | Temperature (°C) | Flow rate (µl/min) | Pumped Volume (µl) | Time (s) |
| --- | --- | --- | --- | --- | --- |
| 1 | Pump Air | 25.2 | 500 | 10 | 0 |
| 2 | Pump hybridization mixture | 38.2 | 500 | 138 | 1200 |
| 3 | Pump Wash Buffer 1 | 25.2 | 625 | 250 | 0 |
| 4 | Pump 3× Saline sodium citrate buffers (SSC) | 25.2 | 625 | 250 | 0 |

Wash Buffer 1: 150 mM NaCl, 15mM sodium citrate, 150 mM HEPES, 0.1% SDS, pH 7.0 @ 25℃.

Surface reverse transcription was performed on the X-bot. Firstly, the program is set to heat the flow cell to 42℃, introduce the RT Reagent, and incubate for 45 minutes. After the reaction, rinse the flow cell, introduce the Exo I Reagent, incubated at 37℃ for 15 minutes, after another rinse, heat the flow cell to 55℃ and rinse with 250 μl volumes of formamide to elute the RNA template. The detailed steps were listed as following.

| Step | Description | Temperature (°C) | Flow rate (µl/min) | Pumped Volume (µl) | Time (s) |
| --- | --- | --- | --- | --- | --- |
| 1 | Take out the reagent kit and change reagent |  |  |  |  |
| 2 | Pump Air | 25.2 | 500 | 10 | 0 |
| 3 | Pump RT Reagent | 43.6 | 500 | 138 | 900 |
| 4 | Pump RT Reagent | 43.6 | 500 | 20 | 900 |
| 5 | Pump RT Reagent | 43.6 | 500 | 20 | 900 |
| 6 | Pump Wash Buffer 1 | 25.2 | 625 | 250 | 0 |
| 7 | Pump 3×SSC | 25.2 | 625 | 250 | 0 |
| 8 | Pump Air | 25.2 | 500 | 10 | 0 |
| 9 | Pump Exo I Reagent | 38.2 | 500 | 148 | 900 |
| 10 | Pump Wash Buffer 1 | 38.2 | 1000 | 250 | 0 |
| 11 | Pump Formamide | 57.6 | 500 | 250 | 60 |
| 12 | Pump Wash Buffer 1 | 57.6 | 1000 | 250 | 0 |
| 13 | Pump 3×SSC | 38.2 | 1000 | 250 | 0 |

RT Reagent (80 µl per flow cell): Mix 16 µl volume of 5 × RT Buffer (250 mM Tris-HCl, 375 mM KCl, 15 mM MgCl<sub>2</sub>, 50 mM DTT, 0.5% Triton X-100, pH 8.5 @ 25°C), 22.8 µl volume of NF-H<sub>2</sub>O, 20 µl volume of 40% Kolliphor® P 188 (prepared in advance, Aladdin, K434430), 3.2 µl volume of dNTPs (each 25 mM), 2 µl volume of Murine RNase Inhibitor, and 16 µl volume M-MLV (Genemind Biosciences Co., Ltd, Shenzhen, China).
Exo I Reagent (50 µl pre flow cell): Mix 25 µl volume of 2 × Reaction Buffer (100 mM KAc, 40 mM Tris-Ac, 20 mM MgAc, pH 7.9 @ 25°C), 21.25 µl volume of NF-H<sub>2</sub>O, 2.5 µl volume of Exonuclease I (Exo I ; Genemind Biosciences Co., Ltd, Shenzhen, China), and 1.25 µl volume of Murine RNase Inhibitor.

TUCL and Sequencing Primer Hybridization , Fill-dATP were also performed on the X-bot. Firstly, rU-tailing Reagent was first introduced to the flow cell and incubated at 37℃ for 30 min. After the reaction, each flow cell was rinsed with 250 μl of wash solution 1, then PK Reagent was introduced for 5 min to remove residual TdT, next, each flow cell was rinsed with 250 μl volumes of wash solution 1 and 3 × SSC sequentially. After that, the flow cell was cooled to 25℃, the sequencing adapter was ligated to the 3’ end of the uracil ribonucleoylated 1^st^ cDNA for 60 minutes. After the reaction, each flow cell was rinsed again to remove the unligated sequencing adapters; Then the Sequencing Primer Mix was introduced, and incubated at 37℃ for 20 minutes to hybridize to the sequencing adapter. Finally, the flow cell was warmed to 42℃, Fill-dATP Reagent was introduced, incubated for 10 min, 2∼4 rU added to the 3’ end of 1^st^ cDNA. Excess poly (dT50LNA2) were extended and filled. The steps above can be completed in four hours automatically. The detailed steps were listed as following.

| Step | Description | Temperature<br>(°C) | Flow rate<br>(µl/min) | Pumped<br>Volume (µl) | Time (s) |
| --- | --- | --- | --- | --- | --- |
| 1 | Take out the reagent kit and change reagent |  |  |  |  |
| 2 | Pump Air | 38.2 | 500 | 10 | 0 |
| 3 | Pump rU-tailing Reagent | 38.2 | 500 | 138 | 600 |
| 4 | Pump rU-tailing Reagent | 38.2 | 500 | 20 | 600 |
| 5 | Pump rU-tailing Reagent | 38.2 | 500 | 20 | 600 |
| 6 | Pump Wash Buffer 1 | 38.2 | 1000 | 250 | 0 |
| 7 | Pump PK Reagent | 57.6 | 500 | 138 | 150 |
| 8 | Pump PK Reagent | 57.6 | 500 | 40 | 150 |
| 9 | Pump Wash Buffer 1 | 25.2 | 1000 | 250 | 0 |
| 10 | Pump 3×SSC | 25.2 | 1000 | 250 | 0 |
| 11 | Take out the reagent kit and change reagent |  |  |  |  |
| 12 | Pump Air | 25.2 | 500 | 10 | 0 |
| 13 | Pump Ligation Reagent | 25.2 | 500 | 134 | 1800 |
| 14 | Pump Ligation Reagent | 25.2 | 500 | 16 | 1800 |
| 15 | Pump Wash Buffer 1 | 25.2 | 1000 | 250 | 0 |
| 16 | Pump Air | 25.2 | 500 | 10 | 0 |
| 17 | Pump Sequencing Primer Mix | 38.2 | 500 | 138 | 1200 |
| 18 | Pump Wash Buffer 1 | 25.2 | 1000 | 250 | 0 |
| 19 | Pump 3×SSC | 25.2 | 1000 | 250 | 0 |
| 20 | Pump Air | 25.2 | 500 | 10 | 0 |
| 21 | Pump Fill-dATP Reagent | 43.6 | 500 | 138 | 300 |
| 22 | Pump Fill-dATP Reagent | 43.6 | 500 | 40 | 300 |
| 23 | Pump Wash Buffer 1 | 25.2 | 1000 | 250 | 0 |
| 24 | Pump 3×SSC | 25.2 | 1000 | 250 | 0 |

rU-tailing Reagent (80 µl pre flow cell): 40 µl volume of 2 × Reaction Buffer was mixed well with 15.2 µl volume of NF-H<sub>2</sub>O, 8 µl volume of CoCl<sub>2</sub> (2.5 mM; NEB, cat. No. B0252SVIAL), 0.8 µl volume of UTP (100mM; Yeasen Bio, cat. no. 10131ES03), and 16 µl volume of TdT (Genemind Biosciences Co., Ltd, Shenzhen, China).
PK Reagent (80 µl pre flow cell): 0.4 µl volume of Proteinase K (20 mg/ml; Sangon Bio, cat. no. B600452-0001) with 79.6 µl volume of PK Buffer (0.1 M Tris-HCl, 0.2 M NaCl, 0.1% SDS, 0.01 M EDTA, pH 8.0 @25°C) to mix well.
Ligation Reagent (50 µl pre flow cell): take 5 µl volume of 10 × T4 RNA Ligase Reaction Buffer (500 mM Tris-HCl, 100 mM MgCl<sub>2</sub>, 10 mM DTT, pH 7.5 @ 25°C) with 21 µl volume of NF-H<sub>2</sub>O, 20 µl volume of 50% PEG8000 (NEB, cat. No. B1004SVIAL), 0.5 µl volume of ATP
(10 mM; NEB, cat. No. P0756S), 1 µl volume of sequencing adapter (33nt\_5'p\_ddC, dissolved and diluted to 50 µM with NF-H<sub>2</sub>O), 2.5 Mix well with a µl volume of T4 RNA Ligase 1 (T4Rnl1, 10 U/µl; NEB, cat. No. M0204L).
Sequencing Primer Mix (50 µl pre flow cell): take 0.5 µl volume of sequencing primers (33 merSP, dissolved and diluted to 10 µM in advance with NF-H<sub>2</sub>O) and mix well with 18.25 µl volume of NF-H<sub>2</sub>O, 31.25 µl volume of hybridization solution V2.
Fill-dATP Reagent (80 µl pre flow cell): 16 µl volume of 5 × RT Buffer was mixed with 47.2 µl of NF-H<sub>2</sub>O, 0.8 µl of dATP (1 mM), and 16 µl of M-MLV.

Sequencing was used the GenoCare sequencing reagent kit and performed on GenoCare1600 to generate 15∼ 40 bp single-end reads (8).

### Data Processing

Low-quality reads were removed using Q values and dustmasker (26). We also removed low-complexity reads, which often result from non-specifically absorbed fluorescent nucleotides. Cutadapt was employed to trim terminal poly-A tails, retaining only reads of 15-30 nt in length. A total of 18 million reads, with a maximum error rate of 10%, were selected for alignment with six miRNA reference sequences. miRNA expression levels were standardized using CPM. For further details on the shortest read length set at 15 nt and the maximum error rate of 10%, refer to the methods section in the supporting information.

### rU-tailing and base bias Optimization

Take 5 μl of 2× Reaction Buffer and mix with 0.6 μl of NF-H2O, 1 μl of CoCl2, 1 μl of UTP (diluted to concentrations ranging from 0.04 mM to 10 mM), 0.4 μl of ssDNA (22 nt-ssDNA, previously dissolved and diluted to 100 μM with NF-H2O), and 2 μl of TdT. Incubate the reaction at 37℃ for 30 minutes, followed by a denaturation step at 70℃ for 10 minutes. Add equal volumes of Novex TBE-Urea Sample Buffer (2 × Loading Buffer; Invitrogen, cat. No. LC6876) to the reaction and analyze on a 20% Novex TBE Urea Gel.

To investigate the influence of the 3’-end base on the rU-tailing reaction efficiency, we designed oligo ssDNAs with different 3’ N1 bases (23 nt-ssDNA-N1C, 24 nt-ssDNA-N1A, 25 nt-ssDNA-N1T, 26 nt-ssDNA-N1G).

We also designed the probe D9TRC-3C-3’Cy3, featuring a 3’ end modified with 3’ Cy3 and complementary to the 5’NH2-C12-D9T region on the chip surface, with a 5’ overhang of CCC. For the D9T probe immobilized on the chip, the test was conducted in one flow cell lane with rU-tailing reagent, while negative and non-specific absorption controls were treated with rU-tailing reagent lacking TdT and UTP in separate lanes. Post-reaction, the test and negative control lanes were hybridized with the D9TRC-3C-3’Cy3 probe at 0.1 μM and extended with a polymerase mix containing G-Atto647N nucleotides (from the GenoCare 1600 R Sequencing Kit V1.0). The addition of rU to the 3’ end of the D9T probe creates an overhang, preventing extension, whereas the negative control group can incorporate

G-Atto647N nucleotides normally. The positive control may adsorb a small amount of G-Atto647N. The colocalization of red G-Atto647N and green Cy3 signals was assessed, with the intensities of G-Atto647N (denoted as IntsG1, IntsG2, IntsG3) and Cy3 (denoted as IntsC1, IntsC2) measured separately. The rU-tailing reaction efficiency was calculated using the following formula:

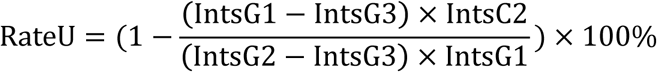

where RateU represents the rU-tailing efficiency, IntsG1 represents the signal intensity of the experimental group after the reaction of G-Atto647N, IntsG2 represents the signal intensity of the image after the negative control group reacts to G-Atto647N, IntsG3 represents the signal intensity of the image after the positive control group reacts to G-Atto647N, IntsC1 represents the signal intensity of the D9TRC-3C-3’Cy3 probe in the experimental group, IntsC2 represents the signal intensity of the D9TRC-3C-3’Cy3 probe in the negative control group.

### ssRNA-DNA Ligation Optimization

The 5’ end of the 22nt-ssDNA-rU3, modified with a hydroxyl group, serves as the acceptor, while the donor adaptor, 33nt-5’p-ddC, features a phosphoric acid modification at the 5’ end and is blocked with ddC at the 3’ end. T4 RNA Ligase 1 (T4Rnl1) is utilized for the ligation.

A range of ATP concentrations from 0.002 mM to 0.6 mM was tested in 1× T4 RNA ligase reaction buffer, with the ssDNA acceptor at a concentration of 1 μM, the donor adaptor at 2 μM, T4Rnl1 ligase at 0.5 U/μl, and PEG8000 at a concentration of 20%. The reaction was incubated at 25℃ for 60 minutes and then terminated by incubating at 65℃ for 20 minutes. The reaction products were mixed with an equal volume of Novex TBE-Urea sample buffer and analyzed using a 20% Novex TBE Urea Gel.

We performed the TUCL reaction was performed as described in section “***TUCL-Seq Library Preparation and Sequencing***”. After the reaction, 0.1 μM of 33nt_SP_5’Cy5 was hybridized in Hybridization Solution V2 at 37℃ for 20 minutes.

Post-hybridization, the samples were imaged using the GenoCare 1600 sequencer. The ssDNA surface ligation efficiency was determined by counting the number of Cy3 and Cy5 fluorescence spots and calculating the ratio of Cy5 to Cy3 spots.

### Surface Superfluity Probes Removal

The poly (dT50LNA2) primer was modified on the surface of the SMS flow cell according to the previous study (25). The X-bot was programmed to deliver Exo I reagent through the flow cell, with treatment times and repetitions set at 15 minutes for 1 to 4 times, and a negative control group treated with reaction buffer. Post-treatment, 0.1 μM A30Cy3 was hybridized using hybridization solution V2 at 37℃for 20 minutes. After hybridization and washing, images were captured on the GenoCare1600 sequencer and analyzed using ImageJ 1.50i software (National Institutes of Health, United States). The Cy3 signal intensity was quantified and compared between the experimental and control groups to determine the digestion efficiency of single-stranded probes on the SMS surface.

### Denaturing poly-Acrylamide gel electrophoresis

ssDNA fragments were determined on denaturing poly-Acrylamide gel electrophoresis using 20% Novex TBE Urea Gel in 1 × TBE Buffer (89 mM Tris, 89 mM Boric acid, 2 mM EDTA·2Na). After electrophoresis, the gel was stained with GelRed (Tanon, cat. no. 170-3001) and photographed with GelDoc XR+ Gel Documentation System (Bio-Rad). Quantitation was performed using ImageJ 1.50i software.

## Data Availability

The authors confirm that the data supporting the findings of this study are available within the article. The data sets used and analyzed are available from the corresponding author on reasonable request.

## Acknowledgments

We thanks professor Runsheng Chen in institute of biophysics, Chinese Academy of Sciences, Beijing for his guidance.We also thanks the employees Haitao Dan and Junhai Liu.

## Author contributions

Q. Yan. conceptualization; JC. Fan, Z. Li, and XJ. You, investigation; JC. Fan, methodology and visualization; JC. Fan, H. Jin, and L. Sun, writing – original draft; H. Jin, PW. Xu, and JJ. Luo data curation, software; L. Liu, and Q. Yan, supervision; YS. Han, P. Wu, resources; L. Sun, and Q. Yan, project administration; YF. Liu writing – review & editing; Q. Yan, funding acquisition.

## Funding and additional information

This work was supported by Shenzhen Science and Technology Innovation Commission and Guangdong Provincial Academician Workstation.

## Supporting Information

This article contains supporting information.

---

**Figure S1.**
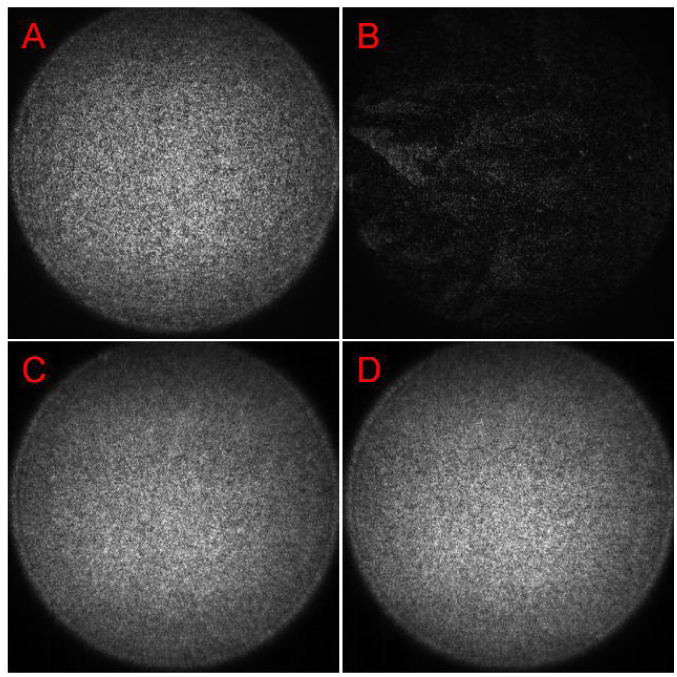
LNA modification enhances the hybridization stability of the miRNA 3’poly(A) tail with the chip surface poly(dT) probe. Poly(dT50) hybridizes with A30RNA-Cy3 before (A) and after (B) the RT reaction, while poly(dT50LNA2) hybridizes with A30RNA-Cy3 before (C) and after (D) the RT reaction.

**Figure S2.**
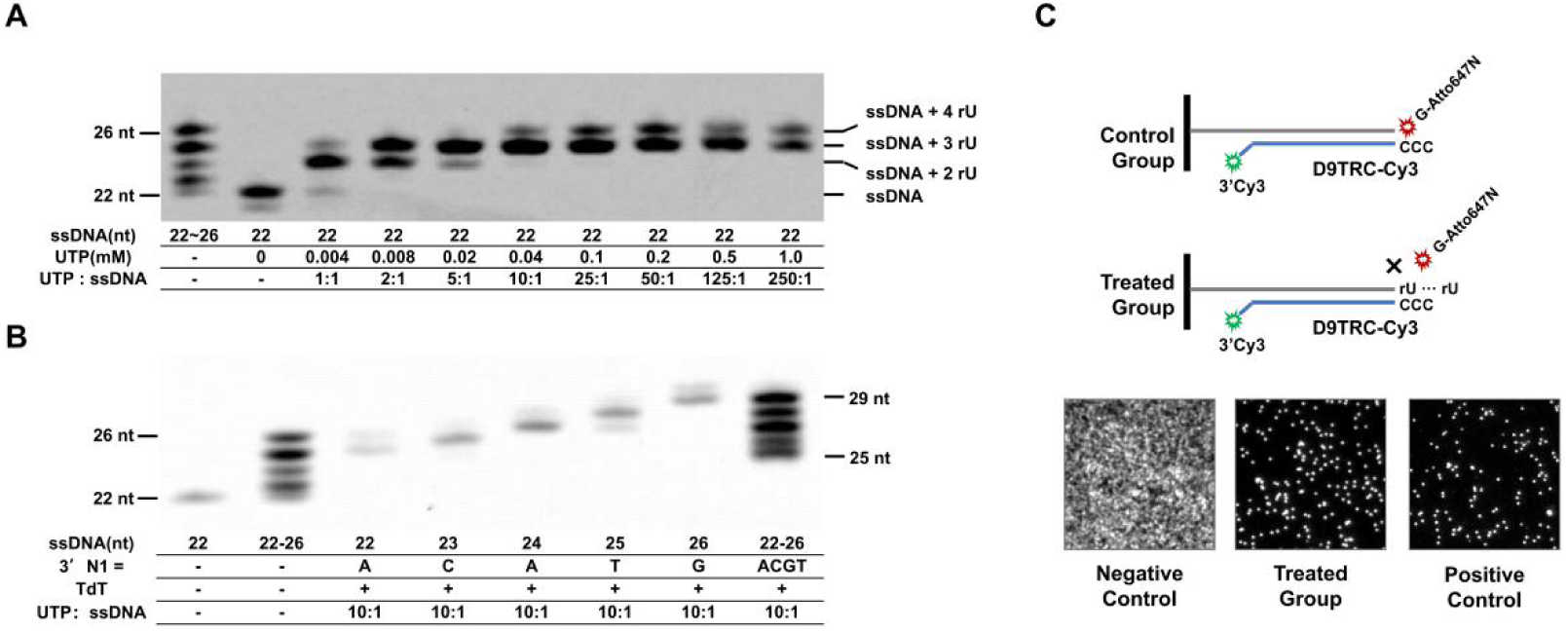
The reaction efficiencyof rU-tailing and 3’ bias. A) Examination of the rU-tailing reaction efficiency at varying molar ratios of UTP to ssDNA, ranging from 0:1 to 250:1; B) Analysis of the rU-tailing reaction efficiency with different 3’ bases on ssDNA; C) Evaluation of the rU-tailing reaction efficiency on the surface of the SMS flow cell.

**Figure S3.**
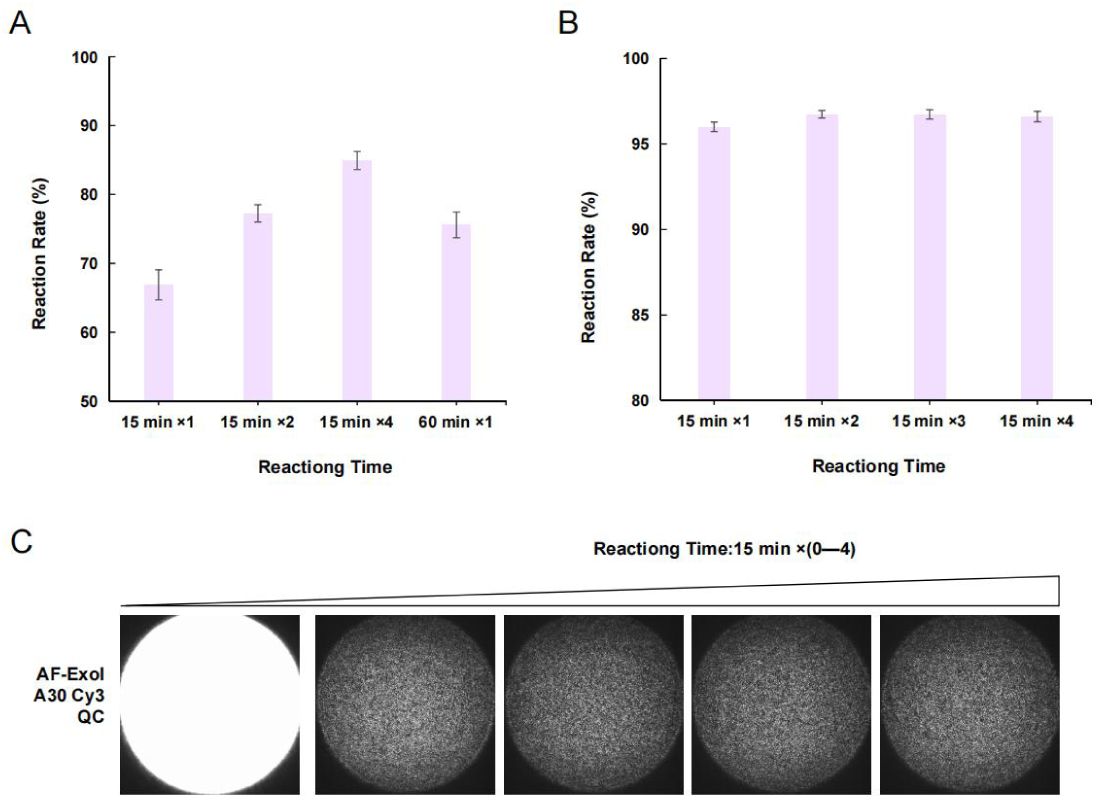
Digestion efficiency of ssDNA probes on the surface of the SMS flow cell. A) Assessment of the digestion efficiency of D9T ssDNA probes is assessed with varying reaction times: 15 minutes (performed once, twice, and four times) and 60 minutes (performed once); B) Assessment of the digestion efficiency of poly(dT50LNA2) ssDNA probes is evaluated with a reaction time gradient, ranging from 15 minutes (reaction 1) to 60 minutes (reaction 4); C) Representative images of the digestion efficiency of poly(dT50LNA2) ssDNA probes is evaluated with a reaction time gradient, ranging from 15 minutes (reaction 1) to 105 minutes (reaction 7), with the leftmost image serving as a negative control, indicating no reaction; D) A schematic representation of the single-stranded configuration of ssDNA probes on the surface of the SMS flow cell.

